# An Open-Set Semi-Supervised Multi-Task Learning Framework for Context Classification in Biomedical Texts

**DOI:** 10.1101/2024.07.22.604491

**Authors:** Difei Tang, Thomas Yu Chow Tam, Haomiao Luo, Cheryl A. Telmer, Natasa Miskov-Zivanov

## Abstract

**Objective:** In biomedical research, knowledge about the relationships between entities, including genes, proteins, and drugs, is vital for unraveling the complexities of biological processes and intracellular pathway mechanisms. Natural language processing (NLP) and text mining methods have shown great success in biomedical relation extraction (RE). However, extracted relations often lack contextual information like cell type, cell line, and intracellular location, which are crucial components of biological knowledge. Previous studies have treated this problem as a post hoc context-relation association task, which is limited by the absence of a golden standard corpus, leading to error propagation and decreased model performance. To address these challenges, we created CELESTA (Context Extraction through LEarning with Semi-supervised multi-Task Architecture), a framework for biomedical context classification, applicable to both open-set and close-set scenarios.

**Methods:** To capture the inherent relationships between biomedical relations and their associated contexts, we designed a multi-task learning (MTL) architecture that seamlessly integrates with the semi-supervised learning (SSL) strategies during training. Our framework addresses the challenges caused by the lack of labeled data by assuming that the unlabeled data contain both in-distribution (ID) and out-of-distribution (OOD) data points. Further, we created a large-scale dataset consisting of five context classification tasks by curating two large Biological Expression Language (BEL) corpora and annotating them with our new entity span annotation method. We developed an OOD detector to distinguish between ID and OOD instances within the unlabeled data. Additionally, we utilized the data augmentation method combined with an external database to enrich our dataset, providing exclusive features for models during training process.

**Results:** We conducted extensive experiments on the dataset, demonstrating the effectiveness of the proposed framework in significantly improving context classification and extracting contextual information with high accuracy. The newly created dataset and code used for this work are publicly available on GitHub (https://github.com/pitt-miskov-zivanov-lab/CELESTA).

## 1 Introduction

Relationships between biological entities, such as genes, proteins, and chemicals, play a fundamental role in driving biological processes. The rapid growth in the amount of biomedical literature has created both opportunities and challenges in studying these complex systems. To address this, natural language processing (NLP) and text-mining tools have been developed to efficiently extract information about relationships between biological entities from large volumes of biomedical texts. Relation extraction (RE) in the biomedical domain, the process of identifying relationships between biological entities in literature, has gained significant attention as an important research direction. In response, substantial efforts have been dedicated to developing extensive biomedical corpora – large collections of texts annotated with biological entities and their relations [1-4]. These annotation tasks typically involve named entity recognition and linking (NER/NEL) to identify the boundaries of biological concepts and link them to curated database entries, followed by RE to determine relations between the identified entities [4]. However, a common issue with biomedical RE is the lack of the ability to identify relevant biological context. Contextual details such as cell type, cell line, and cellular location are crucial for interpreting relations in the biomedical domain, as they provide the necessary information to understand the specific biological relevance, given that interactions can vary depending on the context. For example, protein-protein interactions (PPIs) observed in one cell type may be absent in another due to different regulatory mechanisms, and consequently, differential protein expression. Similarly, gene-disease associations are specific to tissues or organs [5]. A transcription factor’s interaction with another protein might have significant regulatory consequences in the nucleus but be functionally distinct if this interaction occurs in the cytoplasm.

Despite the recognized importance of contextual information, several limitations and obstacles persist in developing effective methods to identify the context of extracted relations. A major limitation is the lack of a golden annotated corpus for this problem, hindering the development of text mining tools, especially deep learning methods that currently rely heavily on manual annotations for training. Furthermore, due to the intricacies of natural language, contextual information may not be explicitly stated in the evidence where biomedical relations are identified, exacerbating the challenges associated with accurate context identification within biomedical texts.

Previous research [6-10] has primarily focused on identifying context words that are explicitly annotated as entities and associated with important concepts such as biomedical events or entities. These studies are limited in scope, recognizing only a narrow range of context types. To address this issue, recent work has employed the pipeline approach, splitting context identification into two sub-tasks: context words are first identified, followed by a binary classification to determine whether the context is associated with the relation [11, 12]. In [11], the authors presented a small annotated corpus and framed the problem as a context-event association task. To build the corpus, their approach employed a machine reader to detect the mentions of both contextual information and biomedical events, followed by expert annotations to establish associations between them. The authors then trained various classifiers, such as Logistic Regression and Support Vector Machines, to extract the context-event relations using syntactic and linguistic features. While innovative, the reliance of this approach on traditional NLP tools may lead to error propagation, potentially affecting the accuracy of context information extraction. The approach described in [12] instead attempts to address the problem of associating context with PPIs using intuitive and interpretable features. Specifically, the authors automatically created a silver standard corpus for training classifiers by extracting sentence-level PPIs associated with specific context mentions, including cell types and tissues. However, this corpus is limited as it concentrates solely on the PPIs extracted from relatively simple sentences and considers only two context types. Moreover, their study assumes that only a context term *c* mentioned within the prepositional phrase “in *c*” in the evidence can be included as a positive context-relation pair, disregarding the possibility of context-relation pairs occurring within complex sentence structures.

Considering the aforementioned limitation, the work in [13] frames the task of recognizing a context-relation pair as a text classification task. The authors introduced a multi-modal model by concatenating the token embeddings and graph embeddings to capture the interactions of biomedical relations and contexts. The model is then evaluated on four context classification tasks: cell line, disease, location, and species. However, in the context annotation classification tasks, the proposed model shows marginal improvement over a baseline NLP model trained solely on textual data. Moreover, the overall performance of these models remains relatively low, suggesting that context classification in biomedical texts is a challenging task that requires further research and development to improve the accuracy and robustness of the solutions.

To address the problems of identifying the context associated with relations mentioned above, we propose *CELESTA (Context Extraction through LEarning with Semi-supervised multi-Task Architecture)*, an open-set semi-supervised multi-task learning (OSSL-MTL) framework for biomedical context classification. Unlike the existing methods, our approach harnesses the advantages of multiple techniques and integrates them into a unified framework. Specifically, to capture the inherent relationships between biomedical relations and their associated contexts, we designed a multi-task learning (MTL) architecture by adding two task-specific classification heads on top of a BioBERT (Bidirectional Encoder Representations from Transformers for Biomedical Text Mining) [14] backbone model. Furthermore, we utilized semi-supervised learning (SSL) strategies to leverage abundant unlabeled data during training. We also created a large-scale dataset for biomedical context classification from two existing corpora and refined it through multiple data filtering and grounding procedures. To enhance the quality of this dataset, support the MTL architecture, and reduce the labor in annotating text spans of entities involved in RE, we developed an entity span annotation method that combines deep learning-based NER models and large language models (LLMs). To demonstrate the effectiveness of our context classification approach, we trained the overall framework on the introduced dataset. We conducted extensive experiments on both close-set and open-set scenarios to examine the impact of the out-of-distribution (OOD) data, providing a comprehensive analysis of the proposed framework.

## 2 Related work

### 2.1 Representations of biological relations and their contextual information

Biological relations can be extracted from unstructured text in literature or found in a structured form in databases [15-17] and knowledge bases [18]. Representation formats that we use in this work include statements in BEL (Biological Expression Language) [19] and INDRA (Integrated Network and Dynamical Reasoning Assembler) [20], and interaction lists in BioRECIPE (Biological system Representation for Evaluation, Curation, Interoperability, Preserving, and Execution) [21].

OpenBEL is a framework that integrates with external tools to support BEL implementation and management [22]. BEL represents biological knowledge in a computable form by describing scientific findings as causal and correlative relationships between entities within the relevant context of biological and experimental systems. BEL statements are structured as *subject*-*predicate*(*relation*)-*object* triples: *subject* and *object* represent biological entities (e.g., protein, biological process, or more specifically, kinase) and the *predicate* (*relation)* indicates the type of interaction (e.g., increases, decreases) between the *subject* and the *object* [23]. A BEL statement is also annotated with additional context information (e.g., cell line, disease, anatomy), a citation and the supporting evidence, including the text of varying length from the cited paper, from sentence fragments to entire paragraphs. We will use the BEL Small and Large corpora from OpenBEL. The Large corpus is the largest public BEL knowledge base, containing over 16,000 publications and 80,000 statements, while the Small corpus includes over 2500 statements from 57 publications. Three examples from the BEL Small corpus are shown in Figure 1(a).

**Figure 1:**
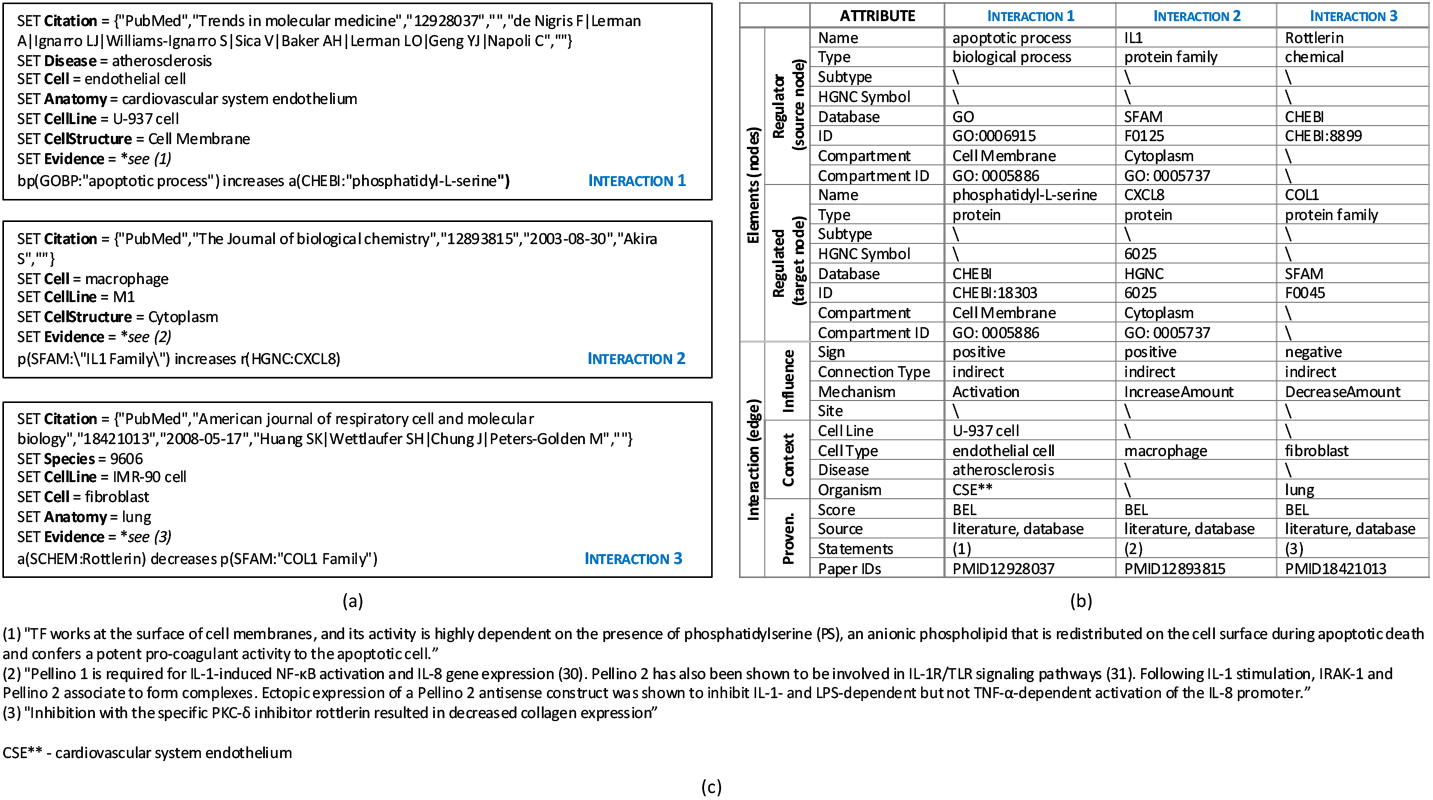
Three interaction examples from the BEL Small corpus and the corresponding representation in the BioRECIPE format. (**a**) Three BEL statements from the BEL corpora, presented in the *BEL script* format. (top box) The *subject* (Regulator in BioRECIPE) of Interaction 1 is a biological process (bp) “apoptotic process”, the *object* (Regulated in BioRECIPE) is an abundance (a) of the chemical “phosphatidyl-L-serine” with the *predicate* (Influence in BioRECIPE) being increases. The information about grounding to the GO and CHEBI databases is also included in both formats. Additional annotations include the Citation (Paper IDs in BioRECIPE) “PMID: 12928037” and the contextual information consisting of Anatomy (Organism in BioRECIPE) “cardiovascular system endothelium”, CellStructure (Compartment in BioRECIPE) “cell membrane” and the associated Evidence (Statements in BioRECIPE). (**b**) Corresponding interactions in the BioRECIPE format, with all attributes. (**c**) The supporting evidence.

INDRA is a knowledge base integrated with multiple NLP and database resources. The output from INDRA is in the form of structured statements representing mechanistic information about biological relations. For example, Interaction 1 (Figure 1(a)) is represented in INDRA as an *activation* statement with a *source* agent (apoptosis) and a *target* agent (phosphatidyl-L-serine). Other annotations are bound in an *evidence* object associated with this statement. Details about INDRA statements, grounding and database references can be found in its documentation [24]. We will utilize some of the functions in INDRA when creating our dataset, as described in Section 3.1.

BioRECIPE [21] is a recently published tabular format that can represent sets of biological interactions, as well as causal and dynamic models, it is compatible with a broad range of used formats (e.g., BEL, INDRA output), and it is simple to read and write by both humans and machines. Directed and signed interactions are written in BioRECIPE with *regulator* (source) and *regulated* (target) *entities* and their *attributes*, rich meta-data, database entries, and *context* and *provenance attributes*. We convert the BEL corpora to INDRA statements and then to an interaction list in the BioRECIPE format to use at the input of our framework. BioRECIPE examples that correspond to BEL statements in Figure 1(a) are shown in Figure 1(b).

### 2.2 Open-set semi-supervised learning in text classification

SSL is a method that incorporates unlabeled data into the training phase to improve learning performance without requiring extensive manual labeling. In NLP, SSL text classification (STC) has proven advantageous in the case when there is an abundance of unlabeled data and labeling is labor-intensive [25]. One of the commonly used techniques of SSL is consistency regularization [26, 27], which aims to improve the robustness and generalization ability of the classification model by enforcing consistent predictions under small changes to the input. For example, Virtual Adversarial Training (VAT) [28], an SSL algorithm based on consistency regularization, improves model’s robustness by training the output distribution to be isotopically smooth around each input through adversarial noise in the direction that most perturbs the output, even in the absence of labels. Another example is Unsupervised Data Augmentation (UDA) [29], which generates augmented data from unlabeled samples by applying augmentation techniques (e.g., back translation, random insertion). UDA calculates the consistency loss based on the difference in predicted outputs, and the total loss is the sum of the supervised loss on labeled texts and the consistency loss on unlabeled texts. The Fixmatch approach [30] combines the advantages of consistency regularization and pseudo-labeling [31] by first generating high-confidence pseudo-labels from weakly-augmented data and then training the model to predict these labels on a strongly-augmented version of the same data.

SSL often makes an unrealistic assumption that all the unlabeled data belong to a known distribution. However, in real-world applications, many unlabeled data may fall into OOD classes that are not defined in the labeled dataset. Therefore, open-set SSL (OSSL) [32, 33] has been proposed to address this challenge by incorporating open-set recognition into SSL, i.e., recognizing the data from unseen classes as outliers. In the OSSL setting, the most straightforward pipeline is to detect and discard samples from unseen classes before performing classification for those from known classes. Various OOD detection methods [34] have been proven effective in identifying outliers in text classification tasks, allowing the language model to focus on the in-distribution (ID) data and improve its classification performance.

### 2.3 Multi-task learning

MTL has shown significant potential in tackling the problem of overfitting and data scarcity in NLP. It has been widely adopted in specialized domains, including biomedical text mining, to leverage the inherent relationships between related tasks through joint training. MTL is typically implemented using two methods: hard and soft parameter sharing [35]. Hard parameter sharing refers to sharing the same model parameters before the task-specific layers, while soft parameter sharing applies constraints to model parameters for different tasks. Recent studies have applied MTL on pre-trained language models (PLMs) such as BERT [36] and its variants for various biomedical text-mining tasks, demonstrating notable improvements over single-task models. For instance, an MTL model with multiple decoders for tasks such as text similarity and RE, NER, and text inference was explored in [37], demonstrating that MTL fine-tuned models outperformed state-of-the-art transformer models trained on single tasks. Furthermore, [38] proposes a transformer-based framework for few-shot biomedical relation extraction, combined with MTL and a knowledge distillation strategy, that outperforms state-of-the-art few-shot learning methods.

## 3 Methods

In Figure 2, we outline the flow diagram of our framework, including the inputs, processing steps, and outputs. Our task is defined as follows: for each context type, given textual evidence in a BioRECIPE interaction, predict the context label from a set of predefined classes. In the following subsections, we first describe our method for constructing the context classification dataset that we used to train and test our framework. Next, we provide details of our framework, which includes the OSSL approach and an MTL architecture, and we define the loss function used in the framework. Finally, we discuss the framework implementation details.

**Figure 2:**
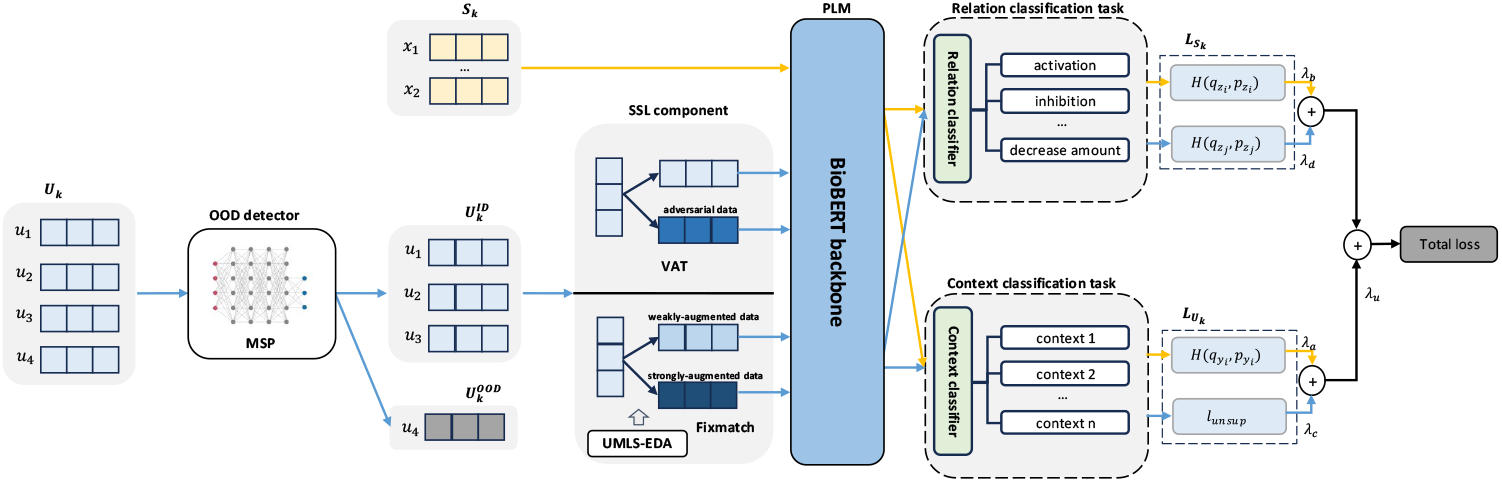
The OSSL-MTL framework for biomedical context classification task. The framework consists of the OOD detector, the BioBERT backbone, relation classifier, and context classifier. Different colored lines indicate data flows for labeled and unlabeled data, with OOD data detected and discarded from the unlabeled set.

### 3.1 Context classification dataset construction

Figure 3 shows the steps we used for creating a large context classification dataset from input corpora. To predict the context from textual evidence without much external biological knowledge, while preserving the precise context-relation association in a corpus, we extended the list of context classification tasks from [13]. Specifically, we retained the “location”, “cell line”, and “disease” context tasks, and removed the “species” task, as obtaining species information from public databases such as UniProt [39] has been proven more efficient than including it as a classification task. We then added two more contexts to our dataset, “cell type” and “organ”, resulting in a total of five context classifications tasks.

**Figure 3:**
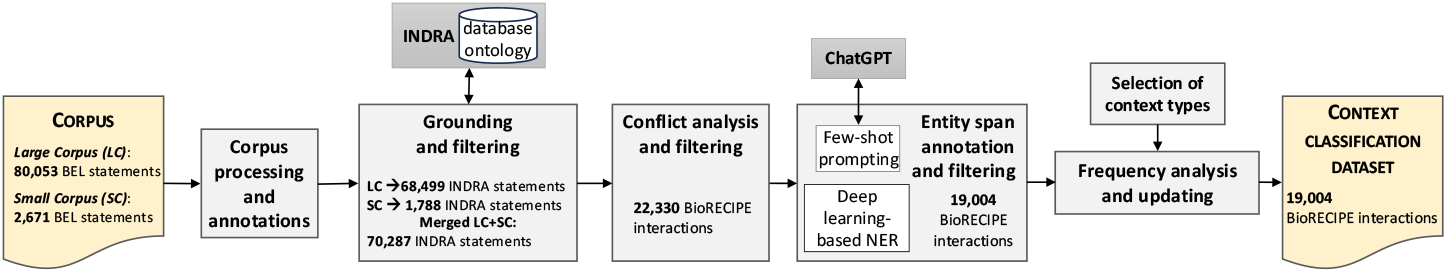
Workflow of constructing our context classification dataset. Evidence (source text) from BEL corpora is filtered and processed by INDRA to ground the regulator (source) entity and regulated (target) entity and the relation with ontology and to link them to database entries. Output from INDRA is investigated for conflicts and further filtered. An entity span annotation method is applied to annotate the text spans with few-shot prompting of ChatGPT and a deep learning-based NER. The final step includes frequency analysis and updating of the dataset to remove the labels with low frequency.

To maintain the high-quality relation and entity annotations in the input corpus and optimize the performance of the context classification algorithm, we filtered and modified the corpus through a series of steps. First, we associated all the annotations (e.g., citations, context and evidence) with BEL statements in the BEL corpus. Next, we grounded relations and entities by utilizing the built-in functions provided by INDRA [20] and removed the statements with relations or entities that could not be mapped to any INDRA ontology or linked to unique identifiers. We processed both Large and Small BEL corpora with these steps, and then concatenated them to generate one larger corpus of INDRA statements (corpora numbers shown in Figure 3). Since we frame the problem as a text classification task, we resolved ambiguities by removing instances with conflicting context labels, ensuring each evidence text has a single context label for its respective classification task. We converted the assembled INDRA statements into a BioRECIPE interaction list.

In biomedical RE tasks, a model is often trained on a corpus to determine the specific type of relationship between pairs of biomedical entities from unstructured text. Hence, annotators must annotate the particular position of biomedical entities in the texts as well as relation types for each entity pair. However, due to the intricate nature of the biomedical ontology and the difficulties of identifying variants of named entities, data annotation for biological relationships requires expertise, resulting in a significant investment of time and effort. Thus, we created a novel Entity Span Annotation (ESA) method that leverages the generalization ability of LLMs and existing pre-trained NER models to annotate entity spans in a corpus.

We first manually annotated five cases as few-shot examples for LLMs (GPT-3.5-turbo) and created a prompt template. We then employed a pre-trained biomedical NER model (en_core_sci_sm) from scispacy [40] to identify the biomedical entities and their corresponding spans within the text. Combining the manually annotated examples with the entity information obtained from the NER model prompted LLMs to automatically annotate the text spans of biomedical entities associated with their relation words. We removed the instances from the corpus that could not be automatically annotated using our ESA method. Such instances typically occurred due to entities being inaccurately annotated in the corpus or inaccurately grounded by INDRA, or their names were too complicated to be recognized.

Next, we analyzed the frequency of different labels used for the five context types in our large corpus. We determined that a minimum frequency of five occurrences in the corpus is an appropriate threshold for class frequency. For each context type, we selected the most frequently occurring classes that satisfy this threshold (Table 1). Since multiple context types can appear within the same statement, we retained instances with higher-frequency labels and replaced less frequent labels—those below our selected class threshold—with an empty string, resulting in our final context classification dataset.

**Table 1:** Context classification tasks (1-5) and their corresponding classes with the highest frequency in the dataset. The relation classification task (6) is an auxiliary task jointly trained within the MTL architecture. For context classification tasks, we include the ontologies to which the contexts are grounded.

| Index | Classification Task | Number of classes | Classes | Ontology |
| --- | --- | --- | --- | --- |
| 1 | Location | 5 | 'cytoplasm', 'cell membrane', 'cell nucleus', 'extracellular matrix', 'extracellular space' | MeSH |
| 2 | Cell Line | 10 | 'HeLa cell', 'MCF7 cell', '293 cell', 'A549 cell', 'NIH-3T3 cell', 'Hep G2 cell', 'U-937 cell', 'THP-1 cell', 'LNCAP cell', 'COS-1 cell' | Cell Ontology |
| 3 | Disease | 9 | 'neuroblastoma', 'breast cancer', 'lung cancer', 'atherosclerosis', 'multiple myeloma', 'leukemia', 'melanoma', 'osteosarcoma', 'lung non-small cell carcinoma' | Disease Ontology |
| 4 | Cell Type | 10 | 'macrophage', 'endothelial cell', 'fibroblast', 'monocyte', 'fat cell', 'smooth muscle cell', 'hepatocyte', 'T cell', 'cardiac muscle cell', 'neuron' | Cell Ontology |
| 5 | Organ | 10 | 'liver', 'lung', 'skeletal muscle tissue', 'endothelium', 'cardiovascular system', 'adipose tissue', 'heart', 'cardiovascular system endothelium', 'blood vessel smooth muscle', 'aorta' | Uberon |
| 6 | Relation | 5 | 'Activation', 'Inhibition', 'IncreaseAmount', 'DecreaseAmount', 'GtpActivation' | N/A |

### 3.2 Creation of the training and test dataset

The majority of data in our dataset have sparse distribution of context labels, with many context attributes being empty in the final BioRECIPE interaction lists. Thus, to ensure that both the training and test sets share a similar distribution, we divided the dataset into six groups based on the presence of labels across the five context types. Each data entry (a row in a BioRECIPE interaction list) belongs to one of these six groups: no labels, one, two, three, four, or five labels across the five context types. As illustrated in Figure 1, Interaction 1 contains five context labels, while both Interaction 2 and 3 lack labels for the cell line and disease contexts. This variation of labeling across interactions creates various combinations of labeled and unlabeled contexts. Once the data was grouped, each group was randomly split into training and test sets using a 70:30 ratio. The training and test sets from all six groups were then concatenated to form the final training and test dataset.

### 3.3 Open-set semi-supervised multi-task learning framework

To effectively capture the context that corresponds to extracted relations while mitigating error propagation, we designed a MTL architecture to train the relation and context classifiers simultaneously. This architecture uses a BioBERT backbone paired with two multi-class classification heads. Furthermore, given the sparsity of context annotations in the dataset and the potential inclusion of OOD examples from pre-defined context classes within the unlabeled texts, we employ OSSL strategies to train our model.

Our framework leverages the unlabeled text set with OOD detection, which is integrated into SSL models with MTL using the two-step pipeline approach from [34, 41]. First, we train an OOD detector using the labeled text set through supervised learning. Next, as illustrated in Figure 2, we apply this detector to filter out the OOD samples from the unlabeled text set. The remaining unlabeled text set and the labeled text set are then fed into the SSL model with MTL for training.

In this study, we use the maximum softmax probability (MSP) method [42] for outlier detection. MSP calculates the softmax probabilities across all classes for each input and identifies data as OOD if the maximum probability (i.e., the model’s confidence on its predicted class) is below a predefined threshold. We focus on SSL strategies based on consistency regularization, where data augmentation is essential for training unlabeled data, as it encourages the model to make consistent predictions on unlabeled data by applying various augmentations. To analyze the impact of data augmentation in our framework, we select the VAT [28] and Fixmatch [30] algorithms as our SSL strategies during training. As Fixmatch requires two types of augmentations (weak and strong), we employ a data augmentation tool named UMLS-EDA [43] that adapt the Easy Data Augmentation (EDA) [44] method to biomedical domains by integrating with the Unified Medical Language System (UMLS) [45] knowledge base. We utilize this tool to generate weakly and strongly augmented data for our unlabeled texts. For weak augmentation, we apply synonym replacement using UMLS-EDA, where biomedical terms are substituted with synonyms from UMLS knowledge base, as we believe that synonym replacement largely retains the syntactic and semantic features of the input text. In contrast, other augmentation methods (e.g., random insertion, deletion) tend to significantly alter the semantics of the text. Therefore, we use these methods for strong augmentation. In the Fixmatch algorithm, the augmented text sets, along with the labeled text set, are then fed into the model during the training phase.

### 3.4 Loss function

The created dataset (Section 3.1 and Figure 3) is divided into five subsets corresponding to different context types. For each context classification task (and context type) *k* ∈ {1, …, 5}, the training set contains both labeled and unlabeled text. In the following, we define the semi-supervised multi-task loss in open-set and close-set scenarios.

Let 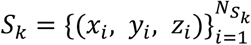 be the labeled text set, where 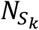 is the number of labeled data of the context type *k*. For each labeled text input *x*_*i*_, it has two corresponding labels, *y*_*i*_ ∈ {0, …, *C*_*k*_ − 1} and *z*_*i*_ ∈ {0, …, *R* − 1}, where *y*_*i*_ denotes the context label (i.e., its corresponding index) and *C*_*k*_ is the number of classes for context type *k* (Table 1), *z*_*i*_ denotes the relation label (i.e., its corresponding index) and *R* is the number of classes for the relation type (Table 1). We denote the unlabeled text set for context type *k* as 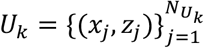, where 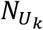 is the number of unlabeled data and *x*_*j*_ are unlabeled text inputs with relation type label *z*_*j*_.

In an open-set scenario, as illustrated in Figure 2, we use the developed OOD detector to split *U*_*k*_ into two subsets 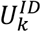 and 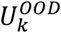, representing the ID and OOD unlabeled data, respectively. During training, a mini-batch is formed by sampling 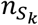 labeled data points from *S*_*k*_ and 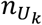 unlabeled data points from either *U*_*k*_ (in a close-set scenario) or 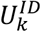 (in an open-set scenario). This leads to the total mini-batch size of 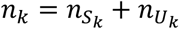 data points for classification task *k*.

The total loss for the backpropagation during training is calculated by combining the losses for the labeled and unlabeled data:

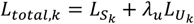

where *λ*_*u*_ is a scalar parameter representing the weight of loss for unlabeled data.

In this work, the loss function follows the standard MTL approach with hard parameter sharing [46], consisting of a context classification loss and a relation classification loss. Within each classification task *k*, we denote the following. The probability distribution for *y*_*i*_ is 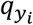, defined as a vector of size *C*_*k*_ with *l*-th component representing the probability for the *l*-th class of the classification type *k*. In the input text set *i*, the probability of the *l*-th component will either be 1, when *y*_*i*_ corresponds to the *l*-th class, or 0, otherwise. Similarly, the probability distribution for *z*_*i*_ is 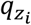, a vector of size *R* whose *l*-th component represents the probability for the *l*-th relation class. Further, we denote 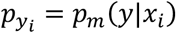 as the model’s predicted probability distribution for context label *y*_*i*_ in the context classification task *k* and 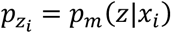 is the predicted probability distribution for relation label *z*_*i*_ in the relation classification task. The loss of the labeled data is then computed as a weighted sum of the two supervised losses for context and relation classification:

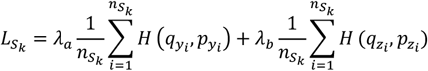

where *H*(*q, p*) is the standard cross-entropy loss function that measures the difference between two probability distributions: the distribution of true labels (actual classes) and the predicted probabilities *p* output by the model [47]; *λ*_*a*_, *λ*_*b*_ are the weight scalars of the context classification and relation classification tasks on the labeled data, respectively.

For the unlabeled data, according to the consistency regularization, when context annotations are absent, the loss comprises unsupervised loss for context classification and a supervised loss for relation classification, as follows:

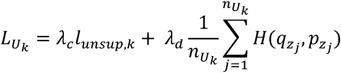

where *λ*_*c*_, *λ*_*d*_ are the weight scalars that determine the relative importance of unsupervised and supervised data, respectively, in the overall learning process; *l*_*unsup,k*_ is the unsupervised loss, which is computed differently and is dependent on the SSL algorithm used (VAT or Fixmatch); 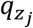 is an input probability distribution for relation label *z*_*j*_ in the unlabeled input text set; 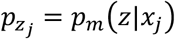 is the predicted probability distribution for relation label *z*_*j*_ in the relation classification task.

The unsupervised loss computed for each context classification task *k*, using the VAT algorithm and the Kullback-Leibler Divergence (KL), is:

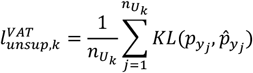

where 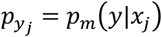 is the model’s predicted probability distribution for context label *y*_*j*_ and 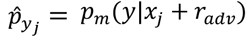 is the model’s predicted distribution when the input *x*_*j*_ is modified by the adversarial perturbation *r*_*adv*_ [28].

In the Fixmatch algorithm, a key distinction from VAT is its reliance on data augmentation techniques to ensure consistency of the unlabeled data and determine the unsupervised loss. In this work, data augmentation is performed using UMLS-EDA, and we denote the weak and robust augmentation as *α* and A, respectively. To obtain the pseudo-label, the class probability distribution 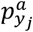 is first computed using the weakly augmented data given unlabeled ID texts such that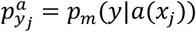. Next, we find 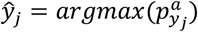 as the pseudo-label and the corresponding probability distribution as 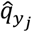. The unsupervised loss is then computed as a cross-entropy loss between pseudo-labels and the output of the strongly augmented *x*_*j*_, which is denoted as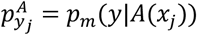. Consequently, the context-specific loss function is as follows:

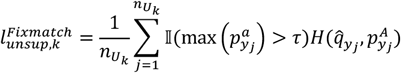

where 𝟙(*condition*) is an indicator function that returns 1 when *condition* is true and 0 otherwise, and *τ* is the probability threshold above which we retain the pseudo-label.

### 3.5 Implementation

The baseline model uses the huggingface [48] package under the default BioBERT parameter settings suggested for sequence classification tasks. We fine-tuned the BioBERT model with 20 epochs as our outlier detector for the MSP method and set the threshold as 0.9. The proposed models are trained with the Adam optimizer [49] for 50 epochs, with a training batch size of eight and a learning rate of 5e-5. Early stopping is applied, and the training is terminated if there is no improvement in the evaluation score for 10 consecutive epochs.

For the MTL settings, we manually tuned the relative weights of two tasks to maximize the predicted score of our model for context classification. All experiments are conducted on our server using a Nvidia a100 GPU.

The predicted results are evaluated on the test set using the conventional metric, micro F1. The Results section exclusively reports the F1 scores of the context classification task, evaluating the models’ efficiency in comprehending and categorizing contexts. The relation classification task provides additional insights into its association with the primary task of context classification.

## 4 Results

In this section, we provide details of our context classification dataset with a summary of the statistics and analysis of the dataset, followed by the evaluation of the performance of the baseline and other SSL/OSSL models with or without MTL architecture on the dataset. We report the F1 scores of classification models with different learning strategies and further analyze the results of each context classification task. Finally, we perform an example analysis of the predicted results on our dataset.

### 4.1 Dataset characteristics and statistics

The context classification dataset contains 19,004 examples from more than 50 biomedical articles adapted from BEL corpora. Table 2 reports the general statistics of our dataset in the open-set scenario across five context types, showing the distribution of the training set, which includes both OOD and ID unlabeled data, as well as labeled data. In the open-set scenario, approximately 10% to 25% of the unlabeled data is identified as OOD data, while in the close-set scenario, the number of unlabeled data for training is the sum of ID and OOD data reported in this table.

**Table 2:** Statistics of context classification dataset on the open-set scenario.

| Dataset | Open-Set # of instance |  |  |  |  |
| --- | --- | --- | --- | --- | --- |
|  | Location | Cell Line | Cell Type | Disease | Organ |
| **Train-unlabeled (OOD)** | 1809 | 3220 | 1124 | 3212 | 1745 |
| **Train-unlabeled (ID)** | 15143 | 13170 | 14072 | 13578 | 13607 |
| **Train-labeled** | 230 | 792 | 1986 | 392 | 1830 |
| **Test** | 103 | 337 | 853 | 168 | 786 |

The data distribution summarized in Figure 4 indicates the number of context attributes annotated across the dataset. Most examples have at most two or three context types labeled, while only 165 instances in the dataset have four context type labels and only four instances are fully annotated with labels for all five context types. The analysis of the top 15 most frequent combinations of any three context types reveals that the predominant cell types are “endothelial cell”, “smooth muscle cell”, and “monocyte”, while the prominent organ types are “cardiovascular system”, “lung” and “blood vessel smooth muscle”.

**Figure 4:**
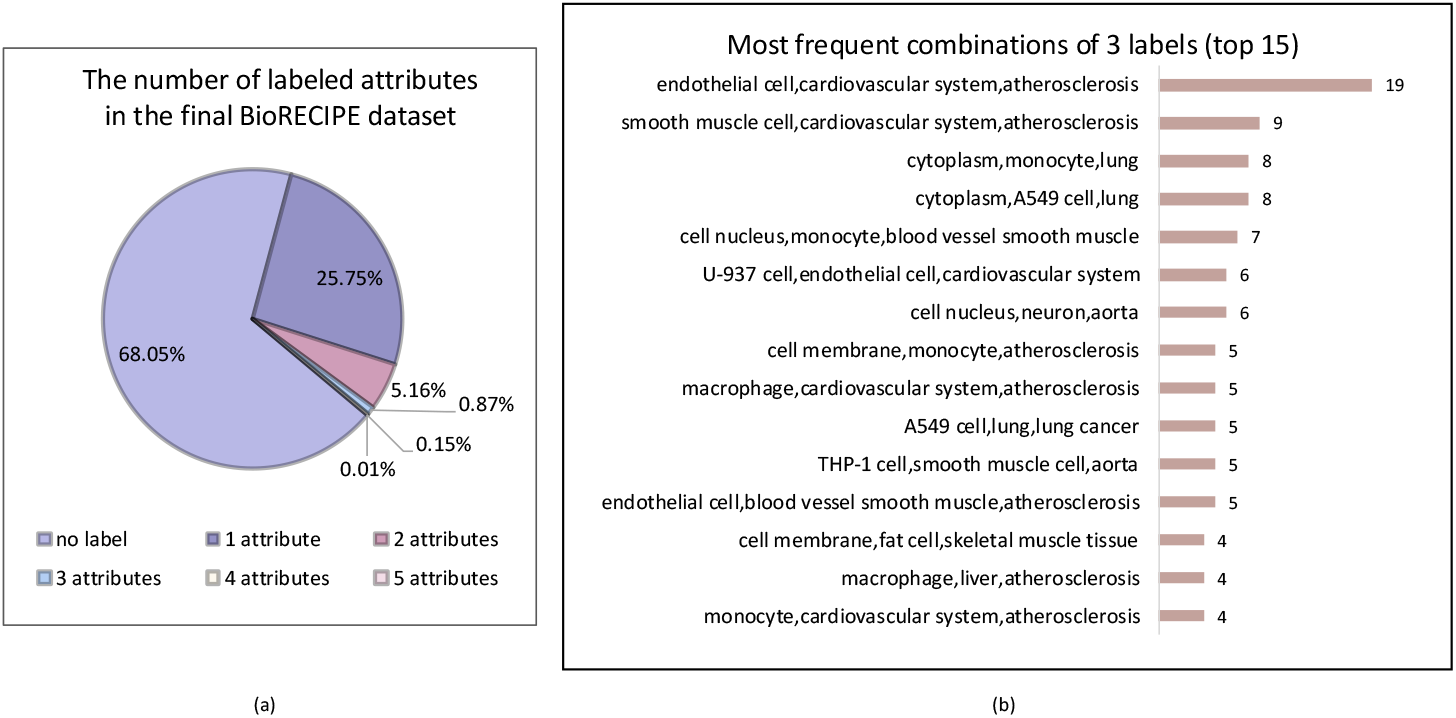
Statistics and characteristic of the context classification dataset. (a) Distribution of labeled context attributes in the final BioRECIPE dataset. (b) Top 15 most frequent combinations of 3 labels in the dataset.

### 4.2 Performance of semi-supervised learning models

Except for the organ classification task, SSL models outperform the baseline by at least 1%, demonstrating their effectiveness in scenarios where supervised data is scarce. The performance of Fixmatch models (Fixmatch, MSP+Fixmatch, MTL+Fixmatch, MTL+MSP+Fixmatch) in Table 3 is relatively inferior to that of the VAT models (VAT, MSP+VAT, MTL+VAT, MTL+MSP+VAT). Interestingly, there is a slight improvement in VAT models with the OOD detection method (MSP+VAT, MTL+MSP+VAT), while this method seems detrimental to Fixmatch models across most classification tasks. Table 3 reveals that most of the unlabeled data are classified as ID data with a threshold of 0.9, which indicates that applying OOD detection before the classification model may not substantially impact our context classification tasks. This also implies that the unlabeled data is of high quality and should be annotated with the same context categories as the labeled data.

**Table 3:** Performance comparison of baseline, MTL, SSL, OSSL, and the proposed models on the context classification dataset.

| Method | Context Classification Task |  |  |  |  |
| --- | --- | --- | --- | --- | --- |
|  | Location | Cell Line | Cell Type | Disease | Organ |
| Baseline | 0.7087 | 0.7211 | 0.7233 | 0.7917 | 0.6921 |
| MTL | 0.7488 | 0.7400 | 0.7238 | 0.7900 | 0.6737 |
| VAT | 0.7312 | 0.7450 | 0.7262 | 0.8037 | 0.6863 |
| Fixmatch | 0.7025 | 0.7438 | 0.7362 | 0.8087 | 0.6850 |
| MSP+VAT | 0.7087 | 0.7488 | 0.7412 | 0.8063 | 0.6875 |
| MSP+Fixmatch | 0.7113 | 0.7262 | 0.7325 | 0.8050 | 0.6850 |
| MTL+VAT | 0.7650 | **0.7562** | **0.7438** | 0.8137 | **0.7113** |
| MTL+Fixmatch | 0.7688 | 0.7325 | 0.7188 | 0.8237 | 0.6913 |
| MTL+MSP+VAT | **0.7775** | 0.7450 | 0.7375 | **0.8287** | 0.6875 |
| MTL+MSP+Fixmatch | 0.7625 | 0.7262 | 0.7250 | 0.8113 | 0.6825 |

### 4.3 Performance of multi-task semi-supervised learning models

Table 3 highlights that the MTL models are superior to SSL and fully supervised models in single-task experiments for location and disease classification. In contrast, SSL models achieve competitive cell line and cell type classification performance. We find significant improvements in MTL+VAT models in both open-set and close-set scenarios (with or without MSP). For example, the MTL+MSP+VAT model performs best in location and disease classification tasks, with the F1 score of 77.75% and 82.87%, respectively. Additionally, the MTL+VAT model surpasses other cell line and cell type classification models. Notably, a slight improvement is observed only for the location and disease classification tasks using the MSP method.

### 4.4 Evaluation

To provide an intuitive understanding of how MTL and SSL improve context classification, we visualize the output embeddings using t-SNE [50] for the baseline model and the best-performing models (based on F1-score) in each context classification task, as shown in Figure 5, with data points from the entire test set. For the location classification task, in Figure 5(a), there is a vague boundary between the embeddings of “cell nucleus” and “cell membrane” for our proposed model, which we attribute to the small size of the test set. For cell type and cell line context types (Figure 5(b-c)), we observe that with MTL and SSL, the label embeddings are well-separated and clustered, whereas the baseline model’s embedding space lacks clear clusters. In the disease classification shown in Figure 5(d), “atherosclerosis” and “lung cancer” classes are forming distinct clusters, suggesting that the proposed model effectively learned their representations. As the improvement observed in the organ classification task is trivial, its t-SNE visualization is not included.

**Figure 5:**
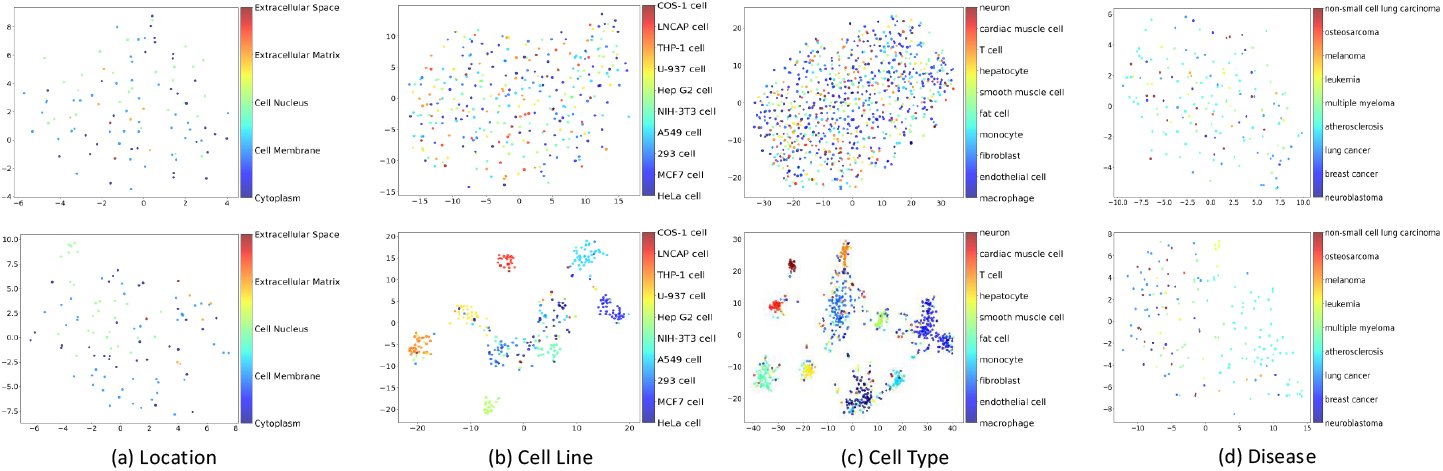
Visualization using t-SNE. The plots illustrate the [CLS] embeddings from the baseline and best-performing models (highest F1 score). Each plot is colored by context labels in the respective classification task, with embeddings assigned to the same cluster sharing the same color. The plots on the top represent the embeddings from the baseline models, and plots on the bottom depict the embeddings generated by best-performing models.

Since it is time-consuming to manually review all the predicted results of our context classification dataset in detail, we conducted a qualitative analysis on five randomly selected instances from the test set, one for each context from the predicted results (Table 4). We observed that if context words were stated explicitly in the input text, most classification models can predict them without any error. However, for more precise context classifications (e.g., non-small cell lung carcinoma, aorta), SSL/OSSL models with MTL architecture outperformed the others, which has shown that MTL could capture the relationship between the interaction and its associated context.

**Table 4:** Analysis of randomly selected examples from the predicted results of the baseline, MTL, SSL, OSSL, and our proposed models on the context classification dataset.

| Context | Input text | Predicted wrong context | Predicted correct context | Models that predict accurately | Reasoning |
| --- | --- | --- | --- | --- | --- |
| **Location** | Deregulation of the pathway results in the aberrant accumulation of <e1>beta-catenin<e1> in the nucleus, often leading to <e2>cancer<e2>. | Cytoplasm, Cell Membrane | Cell Nucleus | MTL+MSP+VAT | Beta-catenin translocates from the cytoplasm to the nucleus |
| **Cell Type** | Protein levels of <e2>RFX5<e2> increase in the nucleus during <e1>IFN-γ<e1> treatment. | T cell, macrophage | fibroblast | MTL+MSP+VAT, MTL+MSP+Fixmatch | Human lung fibroblasts (IMR-90) (Coriell, NJ) and rat fibroblasts (FR) (American Type Culture Collection) |
| **Cell Line** | Exogenously expressed as well as cellular <e1>Hck<e1> and <e2>p73<e2> interact in vivo. In vitro binding assays show that SH3 domain of Hck interacts with p73. Co-expression of p73 with Hck or c-Src in mammalian cells resulted in tyrosine | MCF7 cell, 293 cell, NIH-3T3 cell | None | None | Annotations are wrong, mammalian cells not chinese hamster |
| **Disease** | After treating NSCLC cells containing active Akt/PKB with <e1>LY294002<e1> for 48 h, typical morphological changes of <e2>apoptosis<e2> were observed, including membrane blebbing, increased refractoriness, and chromatin condensation. | lung cancer | non-small cell lung carcinoma | MTL+VAT, MTL+Fixmatch, MTL+MSP+Fixmatch | Non-small cell lung carcinoma is particular form of lung cancer |
| **Organ** | Possible transcriptional regulators of plaque development and inflammation, such as <e1>P-c-jun<e1> and CEBP-b, were down-regulated in <e2>MIF<e2> antibody-treated mice | cardiovascular system | aorta | MTL+Fixmatch, MTL+MSP+Fixmatch | Aorta is a part of the cardiovascular system |

Wrong predictions were due to inaccurate annotations in the original BEL corpora, ambiguous class overlap, class imbalance, and special events such as translocation. Additionally, in some cases the interactions were not unique to any specific cell type or organ, thus the provenance for these contexts is needed to analyze the causes of prediction error, as the models struggled to find answers from the limited input text.

## 5 Discussion

Biomedical relations are inextricably linked to the contexts they are observed in, and therefore, we propose here a solution for identifying biomedical contexts associated with entity relations that are described in biomedical literature. There are several advantages of our OSSL-MTL framework, as discussed in the following.

First, we have created a large context classification dataset by integrating two existing corpora. To remove noise and retain the most accurate annotations, we applied a series of filtering and grounding measures on the dataset. We also assigned a span annotation task to LLMs, which reduces the need of manual annotations and is found to be generally accurate through manual verification. Additionally, we utilized the data augmentation method combined with an external database to enrich our dataset, providing exclusive features for models during training process. The construction of this dataset benefits from data reuse and a set of data processing and automatic annotation methods.

Second, considering the characteristics of our dataset, we designed a MTL architecture to capture the inherent relationship between identified relation and its context. This approach has proven to be effective, as it consistently improves the performance of language models on each context classification task. We noticed that the dataset contains a majority of unlabeled data, suggesting that many of these instances may have corresponding contexts but have not been manually annotated from the original corpora. Thus, we employed SSL techniques to ensure that the abundant unlabeled data can be incorporated in model training. Moreover, we used an outlier detector to filter the OOD data from unlabeled data as not all unlabeled contexts fall within our predefined categories.

This work demonstrates strong adaptive performance in extracting contextual information written in unstructured natural language of biomedical literature. Through our manual verifications of the prediction results, our proposed models have exhibited highly accurate performance on examples where context words are explicitly stated and relatively strong accuracy on those lacking such information compared with baseline models.

There are a few limitations of this work that can be explored further in the future. In our dataset, the annotations of each input only contain a single relation and the corresponding context. However, in certain cases, it is more reasonable to annotate the entities involved in a relation with different contexts. For example, cell membrane receptors interact with extracellular ligands to influence cytosolic signaling, while transcription factors can be activated either in the cytoplasm or the nucleus. Each trained model predicts only one context label given a text sequence. As a result, multiple classification models are required to achieve comprehensive context extraction. Moreover, the categories in the organ classification task are partially overlapping, as aorta, heart, blood vessels and endothelial are all parts of the cardiovascular system. The performance of our models on this task is not significantly improved, as we believe the limited parameters of BioBERT model are insufficient to learn such complex language patterns present in the data. Consequently, investigating the potential of LLMs is a promising avenue for future research. LLMs offer the potential to capture the wealth of contextual information contained within millions of scientific publications and may improve the performance with its capacity of understanding long-distance context association.

The proposed framework in this study can be integrated with any RE model, forming a pipeline for context-specific relation extraction, or regarded as a complementary module to identify the associated context which is vital for extracting cellular events.

## 6 Conclusion

This work combines SSL and MTL approaches for context classification in the biomedical domain. We constructed a large-scale biomedical dataset from existing corpora to reduce labor effort, and annotated entity spans with our automatic ESA method to improve the performance of RE. The problem of identifying context associated with biomedical relations is constrained as a context classification task. Thus, we design an MTL architecture to capture the interaction between relation extraction and context classification tasks. Furthermore, we utilize the SSL approach to use labeled and unlabeled data from the dataset.

The experimental results demonstrate that the proposed approach performs better than the baseline model, single MTL model, and SSL models in most tasks. Further, the performance of different models is compared in both open-set and close-set settings, demonstrating that in a fixed domain, the OOD method may negatively impact the model’s performance. Finally, a detailed analysis for each context classification task is conducted by selecting representative examples for manual evaluation. Future work includes adapting the proposed approach to other information extraction tasks in the biomedical domain and exploring more advanced SSL and OOD methods to improve prediction accuracy of the context classification task.

## CRediT authorship contribution statement

Difei Tang: Conceptualization, Methodology, Software, Validation, Formal analysis, Investigation, Writing – Original Draft, Writing – Review & Editing. Thomas Yu Chow Tam: Conceptualization, Software. Haomiao Luo: Visualization. Cheryl A. Telmer: Data Curation, Writing – Review & Editing. Natasa Miskov-Zivanov: Conceptualization, Resources, Writing – Original Draft, Writing – Review & Editing, Visualization, Supervision, Funding acquisition.

## Acknowledgments

This work was funded in part by the NSF EAGER award CCF-2324742. This research was supported in part by the University of Pittsburgh Center for Research Computing, RRID:SCR_022735, through the resources provided. Specifically, this work used the H2P cluster, which is supported by NSF award number OAC-2117681.

